# Utilisation of Oxford Nanopore sequencing to generate six complete gastropod mitochondrial genomes as part of a biodiversity curriculum

**DOI:** 10.1101/2022.03.24.485721

**Authors:** Mattia De Vivo, Hsin-Han Lee, Yu-Sin Huang, Niklas Dreyer, Chia-ling Fong, Felipe Monteiro Gomes de Mattos, Dharmesh Jain, Yung-Hui Victoria Wen, John Karichu Mwihaki, Tzi-Yuan Wang, Ryuji J. Machida, John Wang, Benny K. K. Chan, Isheng Jason Tsai

## Abstract

High-throughput sequencing has enabled genome skimming approaches to produce complete mitochondrial genomes (mitogenomes) for species identification and phylogenomics purposes. In particular, the portable sequencing device from Oxford Nanopore Technologies (ONT) has the potential to facilitate hands-on training from sampling to sequencing and interpretation of mitogenomes. In this study, we present the results from sampling and sequencing six gastropod mitogenomes (*Aplysia argus, Cellana orientalis, Cellana toreuma, Conus ebraeus, Conus miles* and *Tylothais aculeata*) from a graduate level biodiversity course. The students were able to produce mitochondrial genomes from sampling to annotation using existing protocols and programs. Approximately 4Gb of sequence was produced from 15 Flongle and two R9.4 Flowcells, averaging 235Mb and N50=4.4kb per Flowcell. Five of the six 14.1-18kb mitogenomes were circlised containing all 13 core protein coding genes. Additional Illumina sequencing reveal that the ONT assemblies were able to span over highly AT rich sequence in the control region that was otherwise missing in Illumina-assembled mitogenomes, but still contained a base error of one every 70.8-346.7bp with the majority occurring at homopolymer regions. Our findings suggest that ONT are portable and can be used to rapidly produce mitogenomes at low cost and tailored to genomics-based training in biodiversity research.

## Introduction

Species identification is a key process across biological disciplines [1–4]. Currently, species identity is confirmed through combining morphological and molecular information. In animals, the latter most concerns using mitochondrial markers [5], given their presence in high quantities in metazoan cells and elevated rates of molecular evolution [5,6]. Driven by rapidly improving sequencing technologies and lowered per-base sequencing costs, an approach in which a genome is sequenced to low coverage (often ∼1X) called genome skimming is now available for retrieving and assembling complete mitogenomes from animal samples [9,10]. This is particularly suitable with the Oxford Nanopore Technologies (ONT) platform, a third-generation sequencing method that allows long reads to be sequenced with simple setup [11,12]. For species identification, ONT has been successfully used for amplifying mitochondrial genomes for vertebrates [13] and arthropods [14,15]. Portability has been achieved by ONT with its MinION device, making this technology especially attractive for teaching DNA sequencing and assembly [16–18].

Despite the potential to address genome deficiencies in non-model organisms and for comprehensive species delimitation, ONT mitogenome sequencing is yet to be tested across clades in which it would be extremely beneficial. A taxon of particular interest is the phylum Mollusca [19]. It is the second species-rich animal phylum, with around 117,000 described species and an estimated 150,000 undescribed marine ones [20,21] and has critical ecological, cultural and economic importance [22–28]. According to GenBank ([29], last assessed 27th January 2022), there are 845 mitogenomes (sequences from 13,000 bp onward) available for Gastropoda, 604, for Bivalvia, 224 for Cephalopoda, 4 for Scaphopoda, 3 for Monoplacophora, 24 for Polyplacophora and 9 for Aplacophora. These data have played an important role in understanding evolution in molluscan sub-classes [19,30]. Yet, due to considerable size variation, notable rearrangements, gene duplications and losses as well as reported cases of doubly uniparental inheritance in bivalves, molluscs harbor some of the most complex mitochondrial genomes among metazoans [19].

Here, we establish a system for field-based ONT sequencing of gastropod mitogenomics useful for rapid species identification and mitogenome characterization. We developed a graduate-level curriculum class to specifically address challenges associated with ONT sequencing and assembly and report six high-quality mitogenomes of diverse members of Gastropoda. To assess the accuracy of these ONT assemblies, we produced additional Illumina sequences and compared the extent and nature of sequencing errors and their impacts on mis-assemblies.

## Results

### Sampling and morphological identification of six gastropods

In March 2021, eight graduate students took a sampling trip to Dai Bai Sha on Green Island, Taiwan (**Supplementary Figure 1**). Five gastropod species belonging to four families within Gastropoda were collected and morphologically identified (**Table 1**). The students extracted genomic DNA and sequenced it using Flongle flowcells. We noted that prior to the class a sample DJ was collected in Ruifang, Taiwan to test the whole procedure, resulting in a total of six species presented in this study. All six samples required additional ONT sequence coverage before we attempted to assemble putative mitogenomes. A more formal description of the sampling trip and morphological descriptions are described in **Supplementary Info**.

**Table 1.**
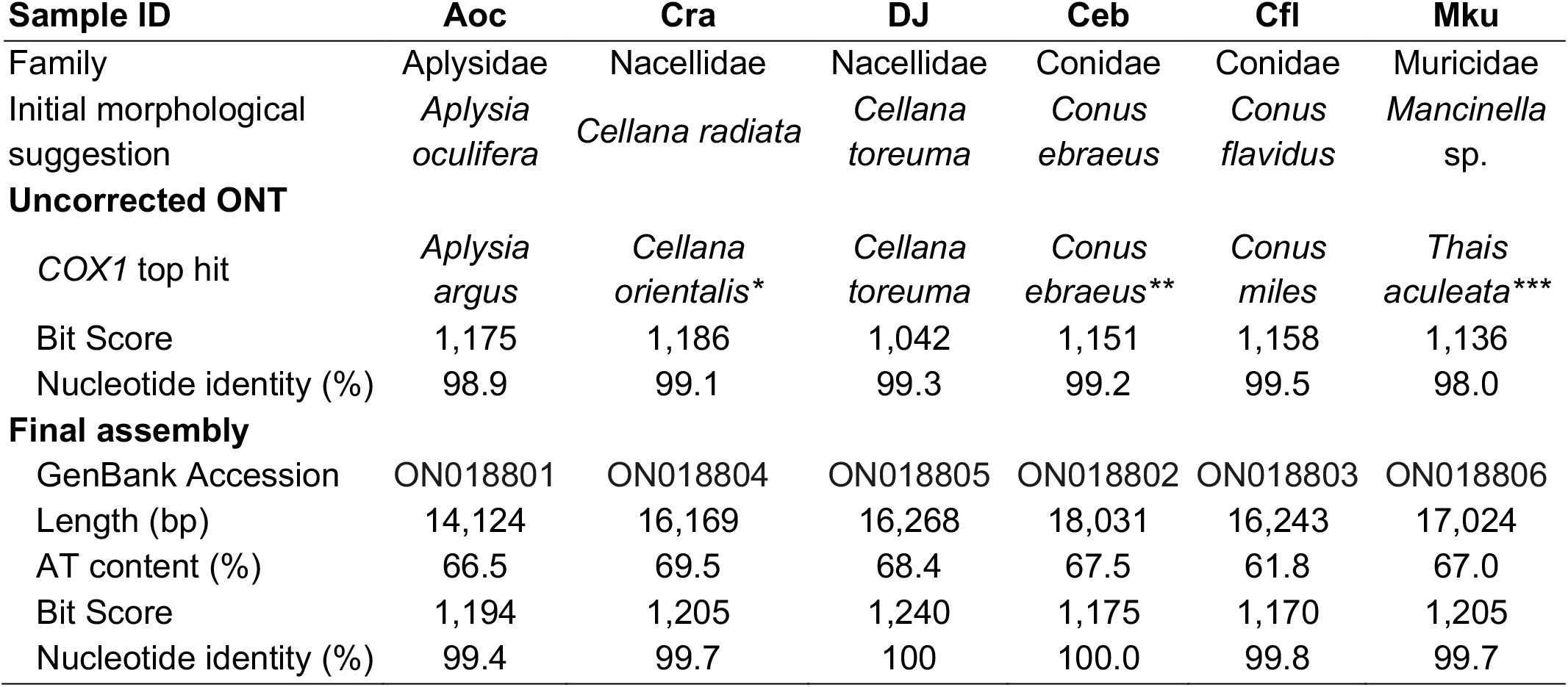
Sample identification (ID) codes, together with original morphological identification and BLASTn results for the whole *COX1* sequence. **\*** Latest species names are provided in the table that were not yet updated in GenBank **\*\*** *Conus cloveri* with 87.2% nucleotide identity was identified as top hit when the full *COX1* sequence was used. We searched instead using Folmer region and identified *C. ebraeus* with much higher nucleotide identity.

| Sample ID | Aoc | Cra | DJ | Ceb | Cfl | Mku |
| --- | --- | --- | --- | --- | --- | --- |
| Family | Aplysidae | Nacellidae | Nacellidae | Conidae | Conidae | Muricidae |
| Initial morphological suggestion | *Aplysia oculifera* | *Cellana radiata* | *Cellana toreuma* | *Conus ebraeus* | *Conus flavidus* | *Mancinella* sp. |
| **Uncorrected ONT** |  |  |  |  |  |  |
| COX1 top hit | *Aplysia argus* | *Cellana orientalis* * | *Cellana toreuma* | *Conus ebraeus* ** | *Conus miles* | *Thais aculeata* *** |
| Bit Score | 1,175 | 1,186 | 1,042 | 1,151 | 1,158 | 1,136 |
| Nucleotide identity (%) | 98.9 | 99.1 | 99.3 | 99.2 | 99.5 | 98.0 |
| **Final assembly** |  |  |  |  |  |  |
| GenBank Accession | ON018801 | ON018804 | ON018805 | ON018802 | ON018803 | ON018806 |
| Length (bp) | 14,124 | 16,169 | 16,268 | 18,031 | 16,243 | 17,024 |
| AT content (%) | 66.5 | 69.5 | 68.4 | 67.5 | 61.8 | 67.0 |
| Bit Score | 1,194 | 1,205 | 1,240 | 1,175 | 1,170 | 1,205 |
| Nucleotide identity (%) | 99.4 | 99.7 | 100 | 100.0 | 99.8 | 99.7 |

### Five out of six circular mitogenomes of gastropods

A total of 17 Flongle and two R9 cells were used to obtain an average of 235Mb of sequence with an average N50 of 4.4kb (**Supplementary Table 1**). Variations in sequencing yield and sequence length differences were attributed to flowcell, library construction, and species specific-characteristics (**Supplementary Table 2**). After filtering for putative mitochondrial reads using the mitogenomes of the most closely related species available in the NCBI database using DIAMOND [31], approximately 10-49X depth of coverage was obtained for each species. Assembly using Flye [32] produced circlised mitogenomes in five out of the six species (**Supplementary Table 3**), confirming that sequencing mitogenomes were achievable in a classroom setting using only sequences from Flongle flowcells and two published programs. Annotation using MitoZ and MITOS [33,34] have shown that five sequences were complete with the presence of 13 protein-coding genes, 22 tRNAs and two rRNAs. An exception was the Cfl sample, which had an incomplete mitogenome lacking the ND5, tRNA^His^ and tRNA^Phe^ genes (**Supplementary Table 3**). At the end of the bioinformatics exercise, students took the annotated COX1 nucleotide sequences and identified the most similar sequences available in the NCBI database via BLASTn or BLASTp. Partial COX1 sequences with 98-99.3% nucleotide identity were obtained in these six samples, providing additional information for species identification (**Table 1**). Three samples (Aoc, Cfl and Mku) had results conflicting with the original morphological identification, which required additional information or phylogenetic analyses to resolve these issues.

### Quantifying the extent of Nanopore errors

Inspection of the annotations from ONT read assemblies revealed the presence of extensive premature stop codons in every annotated protein-coding gene. As a result, only 12-40% of *COX1* query coverage matched a *COX1* homolog in the NCBI nr database using BLASTp (**Supplementary Table 4**). To quantify and correct the extent of errors, we further sequenced the six Gastropoda samples using the Illumina platform. A total of 11.2-47.2X depth of mitochondrial reads were obtained (**Supplementary Table 2**), which were used to *de novo* assemble mitogenomes from Illumina data only as well as to polish the ONT-only assemblies. The consensus quality values (QVs) of Nanopore assemblies were 18.5 to 25.4, corresponding to one base error every 70.8-346.7 bp. Polishing of the ONT-only assemblies using the Illumina sequences resulted in 41-226 modified sites in each species. Comparison of the original to the polished ONT assemblies revealed that the errors were non-random, with single base indels dominating (66.7-85.4%) (**Figure 1A**). Of these, single T and A indels comprised 48% of the total errors presumably because of the high AT composition of mitogenomes (**Table 1**). The majority (71.2%) of errors were located at homopolymer regions (**Supplementary Figure 1**), consistent with previous observations of mitogenome assemblies using ONT technologies [35]. As expected, we observed a positive trend of errors being called with increasing homopolymer length (**Figure 1B**) suggesting it was challenging to basecall precisely in these regions.

**Figure 1:**
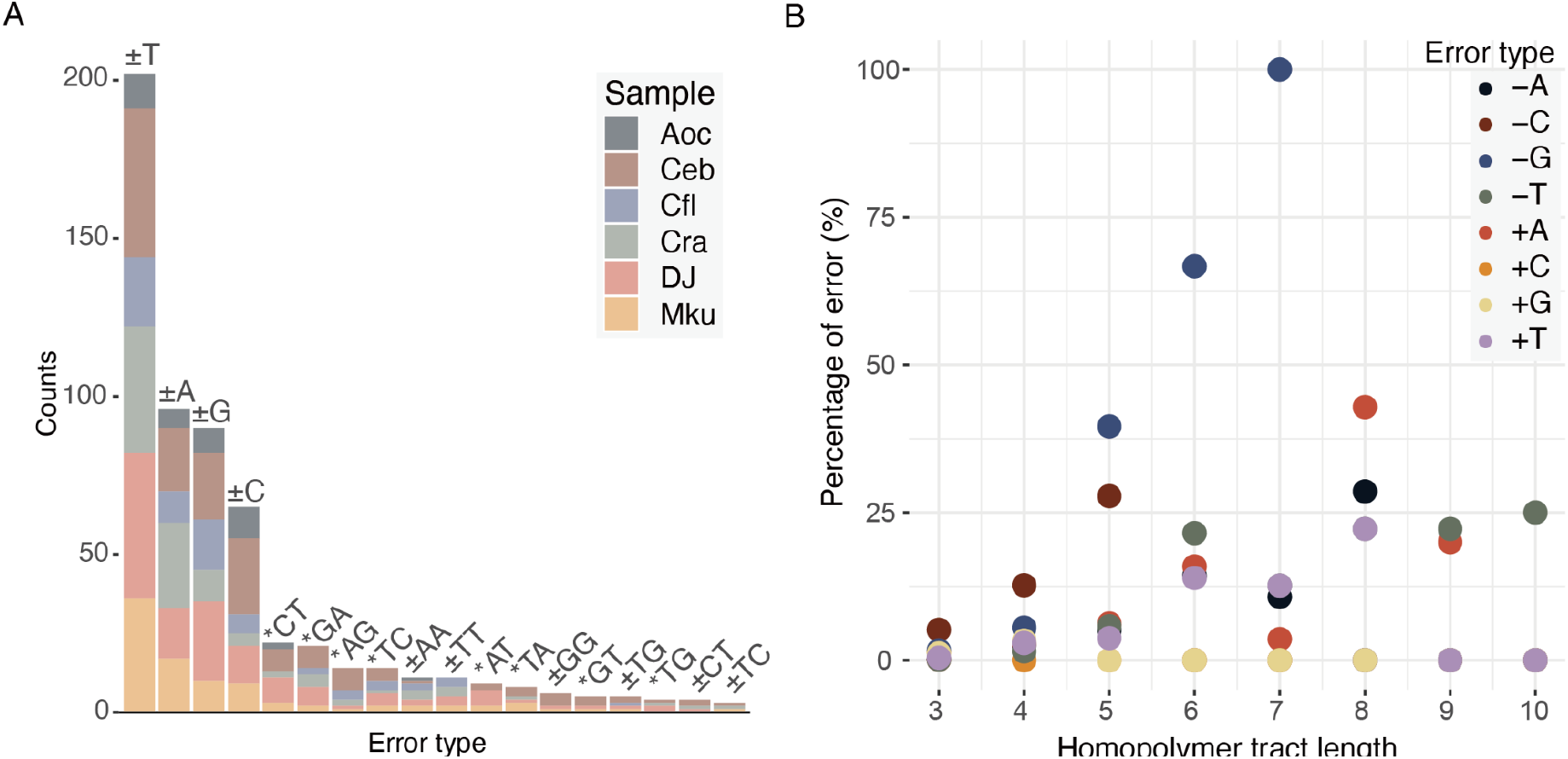
Quantification of ONT errors. (A) Number of INDELs (+/-) and substitutions (*) in ONT assemblies before and after consensus improvement using Illumina reads. Error types that occurred once (n=15) and twice (n=8) were excluded from the plot. (B) Relationship between composition of single-base INDELs and homopolymer length.

We sought to access assembly completeness by comparing the assemblies produced solely from Illumina reads and the polished ONT assembly. In three samples (Aoc, Cra, and DJ), consistent sequences (nucleotide identity 99.9%-100% covering 99.6%-100% of sequence) were observed with both technologies, indicating the assemblies made on these sequences were robust. However, in the Ceb and Mku samples, additional sequences of length 2,169bp and 868bp, respectively were found present only in the ONT assembly (**Figure 2A and Supplementary Figure 2**). The additional sequences are highly AT rich (98.7%; **Figure 2B**) and harbors low Illumina read coverage (**Figure 2C**), consistent with the known property that this technology has difficulties sequencing over regions with highly biased base composition [35]. Despite ONT technology being able to sequence over these regions, a mis-assembly was observed in another sample, Cfl, where one core gene was missing and three were duplicated (**Supplementary Table 3)**. In contrast, the Cfl assembly produced from Illumina reads resulted in all core genes annotated as single copies. The mis-assembly was likely because Cfl had the lowest ONT sequencing N50 (1.2kb) of all the samples despite 27.5X depth of coverage (**Supplementary Table 2**). In comparison, sample DJ produced a circlised assembly with the longest ONT N50 of 8.3kb despite having the lowest depth of mitogenome coverage (11X) amongst samples. For the remainder of the analyses, annotations from polished Nanopore assemblies will be used with the exception of sample Cfl (**Table 2** and **Supplementary Table 5)**. BLASTn results of the polished COX1 sequences showed an increase of 0.3-1.7% nucleotide identity to the same top matched sequences in the uncorrected ONT assemblies (**Table 1**), presumably because the erroneous bases were corrected. As expected, query coverage of the top COX1 hits in BLASTp improved considerably to 99-100% in the final assemblies since they contained no premature stop codons (**Supplementary Table 4**). Together, these results suggest that, currently, a hybrid sequencing approach should be still employed in order to obtain an accurate and complete mitogenome.

**Figure 3:**
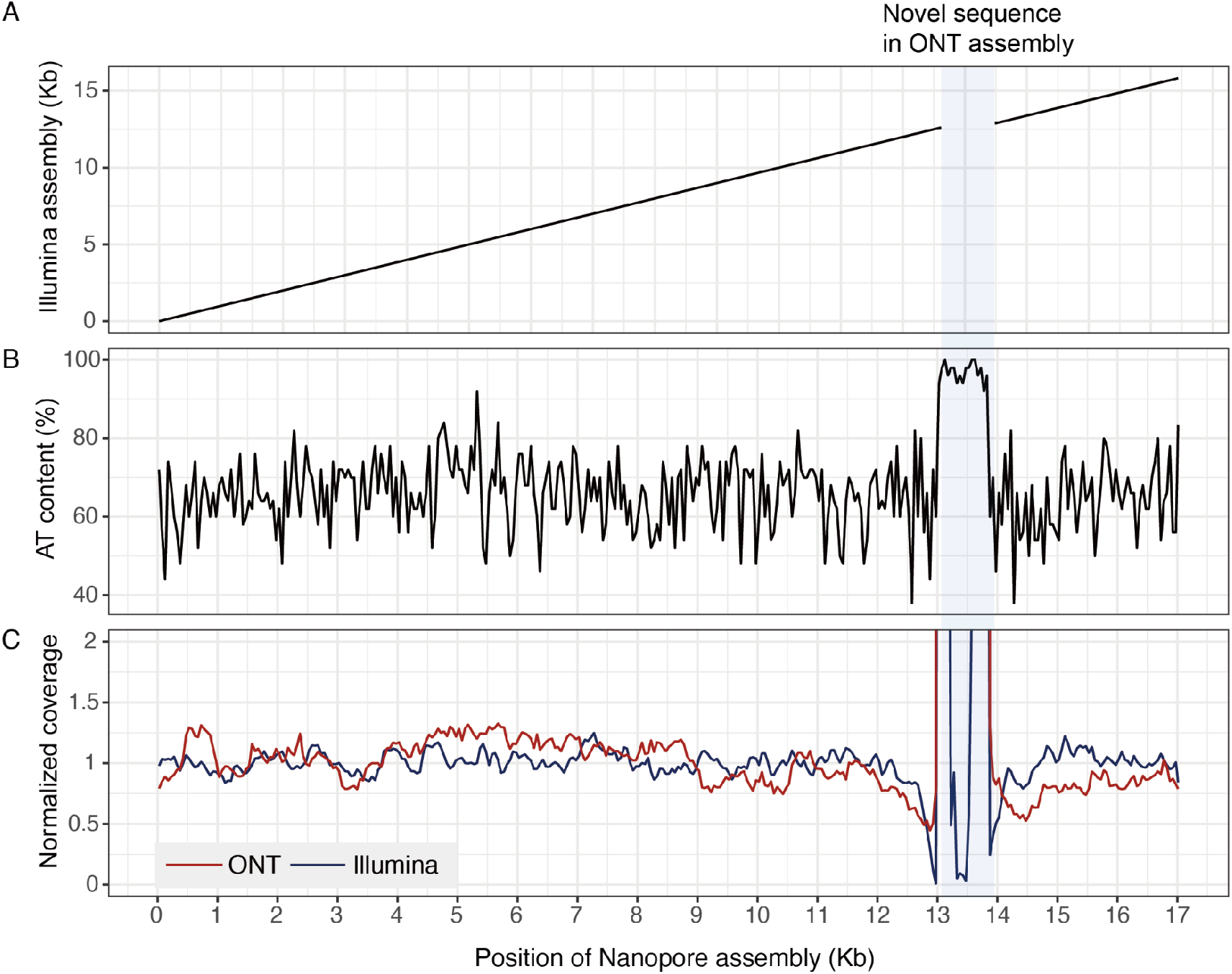
ONT assembly features of sample Mku. (A) Dotplot against Illumina assembly. (B) AT content in 50 bp windows. (C) Nanopore and Illumina read coverage in 50 bp windows.

### Phylogenomics of gastropod mitogenomes

To better resolve species relationships in each family, we constructed a maximum likelihood COX1 phylogeny using nucleotide alignments and mitogenome phylogenies either using concatenated codon alignments of all protein coding genes or coalescence of individual gene phylogenies of representative species (**Supplementary Table 5**). In general, congruence was observed between the COX1 and mitogenome phylogenies, with higher bootstrap support values in the latter (defined here as more nodes with bootstrap>75; **Figure 3; Supplementary Figures 3-7**). With the exception of the DJ sample, all the assemblies reported in this study were the first complete mitogenomes for the designated species.

**Fig 3.**
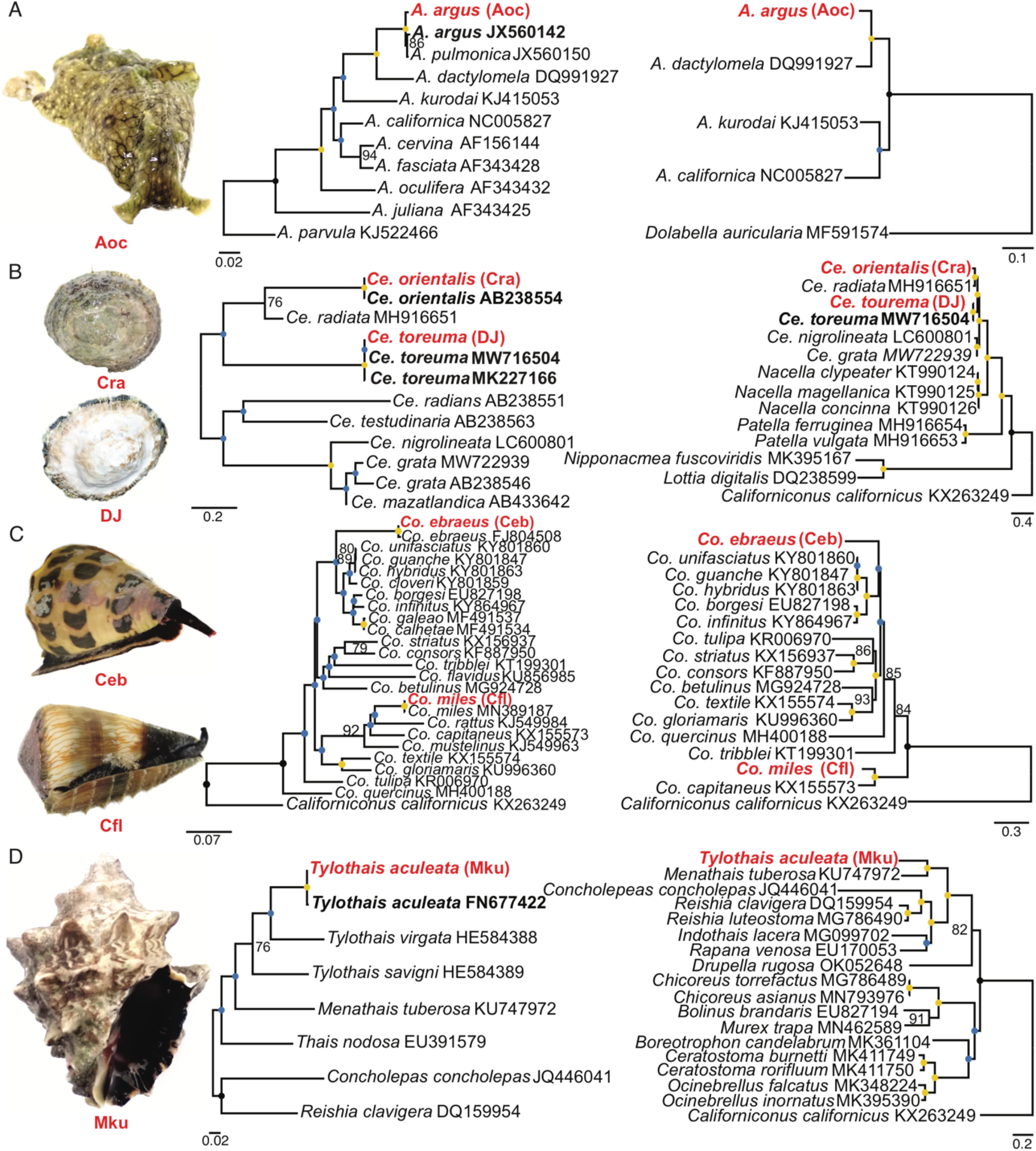
COXI (left) and mitogenome (right) phylogenies from each family. From top to bottom: A) Aplysiidae (with *Aplysia argus*); B) Patellogastropoda (with *Cellana orientalis* and *Cellana toreuma*); C) Conidae (with *Conus ebraeus* and *Conus miles*); and D) Muricidae (with *Tylothais aculeata*). Blue dots represent bootstrap support≤75, yellow ones represent bootstrap support≥95. Values in the middle are written. Red bold tips represent our specimens, black bold ones represent the identified species’ sequences.

Examination of the phylogenetic trees revealed additional information behind four incongruences between initial morphological suggestion and *COX1* top hits. The first was our *Aplysia* species (Aoc, a sea hare), which was originally identified as *A. oculifera* based on the presence of the ring spots alone (**Table 1** and **Figure 3A, Supplementary Info**). We redesignated this sample as *A. argus* (**Figure 3A**) which is the current name used to distinguish the individuals previously recognized as *A. dactylomela* and *A. pulmonifera*’s Indo-Pacific specimens [36], consistent with the clustering in the phylogenies. Second, sample Cra was redesignated as *Cellana orientalis* (**Figure 3B**), which was once regarded as a subspecies of *Ce. radiata* but is now described as an independent species [37]. Third, one of the *Conus* specimens Cfl was initially identified as *Conus flavidus* and redesignated as *Co. miles* (**Figure 3C**). Finally, the murex snail (sample Mku) was tentatively recognized as a species belonging to the genus *Mancinella*,in the taxonomically challenging family Muricidae [38]. We redesignated this sample as *Tylothais aculeata* (**Figure 3D, Supplementary Info**) which was recently erected from *Thalessa* [39] and previously regarded as a *Mancinella* species in Taiwan [40] The Muricidae mitogenome phylogeny was consistent with previous classification, clustering species in the subfamily Rapaninae, Ocenebrinae and Muricinae (**Figure 3D**; [38]).

### Synteny of mitogenomes

The availability of complete mitogenomes allowed us to assess their synteny with sister species and between families. We inspected synteny amongst complete mitogenomes of three Patellogastropoda families (Nacellidae, Patellidae, and Lottidae) and found a general consistency with those from previous studies (**Figure 4 and Supplementary Figure 8**; [41–43]). For example, the most apparent difference, the highly rearranged mitogenomes in Lottidae compared to other Patellogastropoda families, with one large inversion of all protein-coding genes (except COX1 and COX3) between *Nipponacmea fuscoviridis* and *Lottia digitalis* (**Supplementary Figure 15**), was already acknowledged [42]. Interestingly, the control region between tRNA^Phe^ and COX3 typically observed in Gastropoda mitogenomes were much longer in two of our ONT assemblies with the aforementioned novel AT-rich sequences (**Figure 3 and 5**), suggesting hidden diversity present in this region that were previously nearly invisible to Illumina technologies.

**Figure 4.**
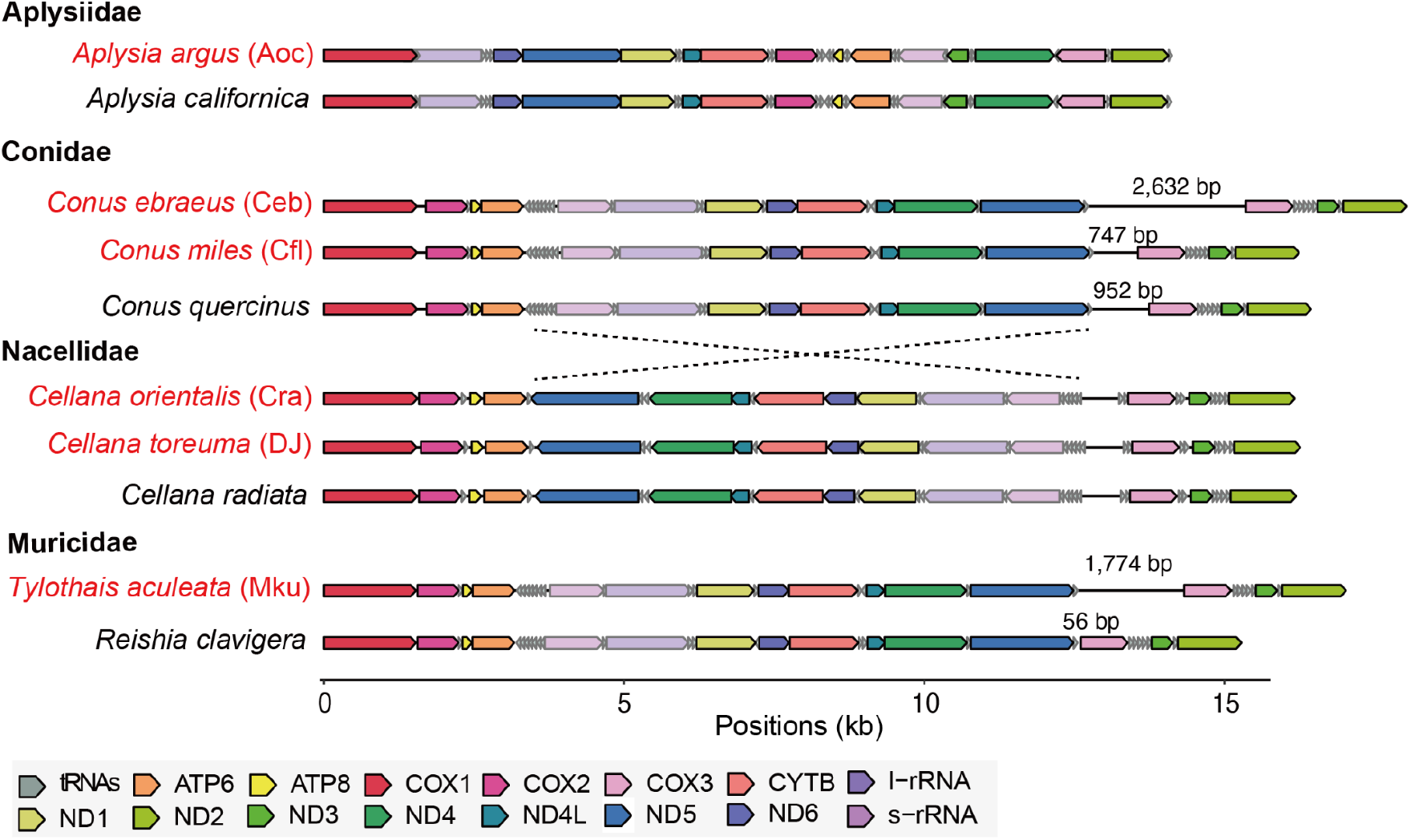
Synteny comparison among our samples and reference mitogenomes. Red labels denote our samples. The lengths of the control region between tRNA^Phe^ and COX3 are shown when very different amongst closely related species.

## Discussion

The primary purpose of this study was to assess whether ONT can be used in a biodiversity curriculum as a reliable tool for generating accurate mitogenomes for expanding resources for the research community. Although multiple assemblies can be constructed and merged in order to achieve greater consensus quality [44,45], we show that closed (i.e., circular) mitogenomes can be achieved with a simple preconstructed bioinformatic pipeline for teaching purposes. This allowed the students to undertake the pipeline and complete the tasks within a typical lecture of three hours. The subsequent polished assemblies can serve as new accurate resources for the research community. Hence, this work highlights that incorporating ONT sequencing in genome skimming approaches holds great potential for exploring and populating sequence databases with the mitogenomes while integrated with educational purposes.

As this was our first attempt to combine ONT with field sampling, sequencing and teaching, we did not target specific taxa and opted for the field sequencing kits that are designed for simple operation and time-efficiency but may compromise the quality of extracted genomic DNA. With students having no *a priori* experience, variations in sequencing yields were anticipated. Despite the prevalence of single base errors, ONT has one advantage over Illumina technology, which is that the sequencing of highly AT-rich sequences is not biased. The novel AT-rich sequences in *C. ebraeus* and *T. aculeata* coincided with the control region amongst published assemblies (**Figure 4**) and implies a re-assessment using different sequencing technologies may be useful. Recently, long read sequencing has corrected errors in at least 100 reference mitogenomes [46]. Given the anticipated increased performance as ONT matures, confirmation and validation with additional ONT sequencing may be built into part of the teaching curriculum to specifically address samples that have suspect control region assemblies.

In conclusion, this study shows that ONT can also be a tool for students to learn how to work with and sequence DNA directly in a field station, thus making it fit as part of a graduate-level class and curriculum in biology and bioinformatics. Although the qualities of the ONT-only assembly can be improved through multiple passes and comparisons of assemblies using different programs, this was not feasible in the teaching context. A hybrid sequencing approach with an initial ONT assembly and polished with Illumina reads is far more straightforward and certainly can be integrated into the curriculum. With continuous improvement in read accuracy and yield in long read technologies, we anticipate one day that new accurate and complete mitogenomes may rapidly populate the Tree of Life across different corners of the world by users ranging from evolutionary biologists to citizen scientists to high school students.

## Material and Methods

### Sampling processing, DNA extraction and sequencing

Sampling of six gastropods by the students is detailed in **Supplementary Info**. The solutions used to extract high-yield genomic DNA for mitogenome sequencing were prepared prior to field sampling and DNA extraction following the manufacturer’s instructions. We used the Quick-DNA™ HMW MagBead Kit (Catalog No. D6060) for DNA extraction and then the DNA samples were stored in a fridge (4ºC) before Nanopore mitogenome sequencing.

For Nanopore long read sequencing, ∼400ng of genomic DNA per sample were used for library construction. Sequencing library was generated using the Field Sequencing kit (SQK-LRK001, Oxford Nanopore Technologies, UK), following the manufacturer’s instructions. 30 μl or 75 μl of the library were loaded into a Flongle or R9.4 flow cells, respectively. Each library was sequenced by a MinION device for 24-48 hours. The ONT FAST5 output files were converted to FASTQ files using Guppy 4.4.2 (https://nanoporetech.com/community) in fast mode with default setting (Oxford Nanopore Technologies, Oxford, UK).

For Illumina short reads sequencing, ∼200 ng DNA per sample was used for the DNA library preparations. Sequencing libraries were generated using TruSeq Nano DNA HT Sample Prep Kit (Illumina USA) following manufacturer’s recommendations and index codes were added to each sample. Briefly, genomic DNA sample was fragmented by sonication to 350 bp. Then DNA fragments were end-polished, size selected, A-tailed, and ligated with the full-length adapter for Illumina sequencing, followed by further PCR amplification. After PCR products were purified (SPRIselect reagent, Beckman), libraries were analyzed for size distribution by Agilent 2100 Bioanalyzer and quantified by Qubit. The DNA libraries were sequenced on the Illumina NovaSeq 6000 platform and 150 bp paired end reads were generated by Genomics BioSci & Tech Co. Illumina reads were trimmed by fastp (ver. 0.22; [47]) with default parameters.

### Assembly and annotation of Gastropod mitogenomes

Amino acid sequences of the complete mitogenomes of sister species to the samples were obtained from NCBI (Sample Aoc: *Aplysia californica* NC005827.1; Ceb and Cfl: *Conus quercinus* NC035007.1; Cra and DJ: *Cellana radiata* MH916651.1; and Mku: *Reishia clavigera* NC010090.1). These sequences served as baits to search for putative mitochondrial sequences using DIAMOND (ver. 0.9.24.125; [31]). An initial assembly was produced from these putative mitochondrial sequences using Flye (ver. 2.8.3; [32]) and served as baits to search for all possible mitochondrial sequences using Minimap2 (ver. 2.24; options: -x map-ont; [48]). A second round of ONT assemblies were produced and further polished using the same set of data by racon (ver. 1.4.11; [49]) for four iterations and medaka (ver. 1.2.0; https://github.com/nanoporetech/medaka). A final round of polishing was conducted using Pilon (ver. 1.22; [50]) with Illumina reads. Assemblies using solely Illumina reads were generated using MitoZ (ver. 2.4-alpha; [33]). Both versions of assemblies were subjected to MitoZ (options: --clade Mollusca) for annotation. The one which had better sequence integrity and gene completeness was selected as the final version. Gene annotations on final assemblies were further curated manually to ensure correctness. Read mappings for long and short reads were performed using Minimap2 (ver. 2.24; options: -x map-ont; [48]) and bwa (ver. 0.7.17; [51]), respectively. Duplicates in Illumina mappings were marked by SAMBLASTER (ver. 0.1.26; [52]). The estimation of read coverage was conducted by Mosdepth (ver. 0.2.5; [53]). The comparison between assemblies was conducted using Minimap2 (options: -x asm5 --cs) and the paf format output was parsed. Part of the pipeline was redesigned as a three-hour lecture available at https://introtogenomics.readthedocs.io/en/latest/emcgs.html and detailed in **Supplementary Info**.

### Phylogenetic and synteny analysis

We used 13 mitochondrial protein-coding sequences to construct trees within the gastropod families Aplysiidae, Conidae, Muricidae and subclass Patellogastropoda. For the reference sequences, we used mitogenomes within family Aplysiidae (5), Conidae (18) and Muricidae (17) and within subclass Patellogastropoda (13) as reference, which were downloaded from GenBank ([29]; last assessed: 18th February 2022). The details of downloaded references are shown in **Supplementary Table S5**. Concatenated and coalescence methods were applied to codon alignments of 13 protein encoding genes in our newly sequenced samples and reference sequences. Sequence alignments for each mitochondrial protein-coding gene was performed using the L-INS-i algorithm in MAFFT 7.487 [54]. We concatenated the genes by using SequenceMatrix [55] and then built Maximum Likelihood phylogenies using ModelTest and RAxML-NG implemented in raxmlGUI [56], with 500 bootstraps replicates. A consensus tree based on coalescencing all individual gene phylogenies were constructed with ASTRAL [57]. The trees were visualised with FigTree 1.4.4 [58]. Gene order of mitogenomes were visualised using the gggenomes package (https://thackl.github.io/gggenomes/).

We downloaded COI sequences from GenBank for checking the species ID. The sequences were chosen according to a BLASTn search [59] with default settings. The alignment was performed with MAFFT 7.471 [54] and trimmed manually while inspecting the alignments under MEGA X 10.1.8 [60]. In total, 646 bp were used for reconstructing the *Aplysia* COX1 tree, 636 for Conidae, and 657 for both Muricidae and *Cellana*. After that, we used ModelTest and RAxML-NG implemented in raxmlGUI [56] for building a Maximum Likelihood phylogeny for each clade, with 500 replicates. If there were issues with scientific names (i.e., synonyms), we decided to use the ones accepted by the World Register of Marine Species [37].

## Authors’ contribution

B.K.K.C., J.W., R.M. and I.J.T. conceived the project. M.D.V., H.-H.L., Y.-S.W., N.D. and I.J.T. analyzed the data and wrote the manuscript with input from others. M.D.V., Y.-S.W., N.D., C.-l.F., F.M.G.d.M., D.J., Y.H.V.W and J.K.M. sampled the specimens, extracted the DNA and performed the initial ONT run. T.-Y.W. supervised the ONT sequencing and sequenced the subsequent runs. Order of first authorship was determined by three rounds of Street Fighter II and King of Fighters ‘97, and the co-first authors have the right to list their name as first in their CV as they contributed equally.

## Acknowledgement

We thank Pei-Chen Tsai, Yao-Feng Tsao and the staff of Green Island Marine Research Station, Marine Science Center (Academia Sinica) for help with specimen collection and field trip logistics, Vanessa Chen (Academia Sinica) for helping with organizing both the class and the trip to Green Island. We kindly acknowledge support from Tzu-Ching Meng (Academia Sinica), who started this first TIGP signature course, Ecology Master Class Taiwan (EMT) and whose support was very important to the completion of this work. This study was supported by the Taiwan International Graduate Program (TIGP) and Biodiversity Research Center, Academia Sinica (Taipei, Taiwan). ND was jointly sponsored by a double-degree graduate grant from TIGP and the Natural History Museum of Denmark. This is EMT paper #2.

