## Supplementary Info for "Utilisation of Oxford Nanopore sequencing to generate six complete gastropod mitochondrial genomes as part of a biodiversity curriculum"

This file includes:

1. Supplementary Text: Extended Results
2. Supplementary Figure 1 to 8

Other supplementary materials for this manuscript includes:

Supplementary Tables are in a merged Excel file.

#### Supplementary Text

##### Field sampling

Prior to the start of a graduate class taught at Taiwan International Graduate Program for Biodiversity (https://tigp-biodiv.biodiv.tw/index.php/emt-tigp-signature-course/), one sample (DJ) was collected and sequenced in Ruifang (25°07'17.8"N, 121°49'19.9"E), Taiwan to test the protocols laid out in this study. All materials were used for this sample, hence no voucher was kept. The students went on and collected five Gastropoda samples from the rocky shores in Da Bai Sha (Green Island, Taiwan, 22.639° N, 121.493° S; WGS84, see **Supplementary Figure 1**) during low tide on 22nd March 2021. Specimens were either collected by hand or with tweezers. The specimens were placed in 2-5 L transparent bottles with sea water and transported back to the lab at the Green Island Marine Research Station, Marine Science Center, Academia Sinica, Taiwan.  Here, they were separated by taxon and kept in 10-20 L aquaria in aerated seawater. The specimens were not fed prior post-sampling processing. After collecting explants/tissue samples for DNA extraction, the remaining parts of the specimens were fixed in 95% ethanol and brought to Academia Sinica Museum of Natural History (sample IDs ASIZM0001713, ASIZM0001714, ASIZM0001715, ASIZM0001718, ASIZM0001719). Each specimen was carefully removed from their aquarium tanks and placed on roughly 20 x 20 aluminum foil pieces. We used sterilized scissors or razor blades to dissect four 25 mg muscle tissues from each specimen of either the subepithelial tissue, muscular foot or heart.

##### Morphological description of gastropods

*Aplysia* is a genus of sea slugs under the order Anaspidea and family Aplysiidae. Sea slug species have highly reduced internal shells and the body of several *Aplysia* species is characterized by ring-like spots over the head and parapodia. Given this, species identification can be challenging if done morphologically, due to the presence of color and marking polymorphism [1]. In particular, *A. argus* resembles *A. oculifera* and *A. dactylomela*, although the latter is not regarded  as occurring in the Indo-Pacific region anymore and the mentioned species can be recognized by their internal shell morphology [1].

*Cellana* is a limpet (superfamily Patelloida) genus belonging to the family Nacellidae, known for having a singular flattened shell  and for being grazers which feed on algae as food [2,3]. *Cellana* is widespread in the Indo-Pacific area [4] and its species were usually recognized by shell morphology, although its high variability made it complex and now usually radular and body characteristics are used instead [2]. As a proof of the complexity of the characters used for species delimitation in this genus, *Ce. orientalis* has a very controversial taxonomy and was once regarded as a subspecies of *Ce. radiata* [5]. The latter is regarded as a species complex [3]. As a possible morphological marker, Powell [6] reports "very distinct radial folds that underlie the normal radial sculpture" in *Ce. orientalis* (although it was still regarded as a subspecies of *Ce. radiata* at that time). Regarding *Ce. toreuma*, it is widely distributed in East Asia. It presents wide variation in sculptural and color characters, although it presents a depressed conical shape and the aperture is usually oval. It has been present in Asia at least since the Middle Pleistocene [7].

The genus *Conus* is a predatory clade, with more than 750 described species [8]. It has thick coiled shell with the whorls enrolled upon themselves and short shell spire. The aperture is narrow and elongated with parallel margins and they also have a needle-like modified radula with a venom gland to attack and paralyze the prey and then engulf it [8–10]. Both the species we sampled feed on polychaetes [9]. From what concerns *Co. ebraeus*, it superficially resembles *Co. judaeus* due to the presence of black botches on the white shell, although their radula and their feeding preferences (*Co. ebraeus* feeds on Eunicidae, while *Co. judaeus* on syllids) are different [11]. *Co. miles*, instead, share some very vague similarity with *Co. capitaneus* but it has brownish lines around the whorls and spire [9].

*Tylothais aculeata* is a murex (family Muricidae) snail, which preys on other intertidal invertebrates [12,13]. The shell is coiled and has shell spines [13]. It was recently erected as a standalone genus, due to the fact the previous genus (*Thalessa*) is now regarded as a junior synonym of *Volema* [13]. It was regarded as a *Mancinella* species in Taiwan faunal checklist [14].

**Design of student bioinformatic class**

Prior to the analysis class, a questionnaire was given to the students to assess their experiences in genome skimming and familiarity with Linux. As approximately half of students had limited or no experiences with Linux, the whole exercise was designed to run under three hours and was conducted by pairs of students with at least one student having some experiences with Linux. The whole exercise is available in at https://introtogenomics.readthedocs.io/en/latest/emcgs.html from assigning putative mitochondrial reads to assessment of assembly metric. As MitoZ required more time to run, the finished results were prerun and available to students who have completed all the exercise for further interpretation of the annotated results.

### Supplementary Figures

**Supplementary Figure 1.** Sampling site of gastropodes. Gastropods were collected from a rocky shore at Da Bai Sha, Green Island, Taiwan (22.639° N, 121.493° S) and brought back to the Green Island Marine Research Station for further processing.

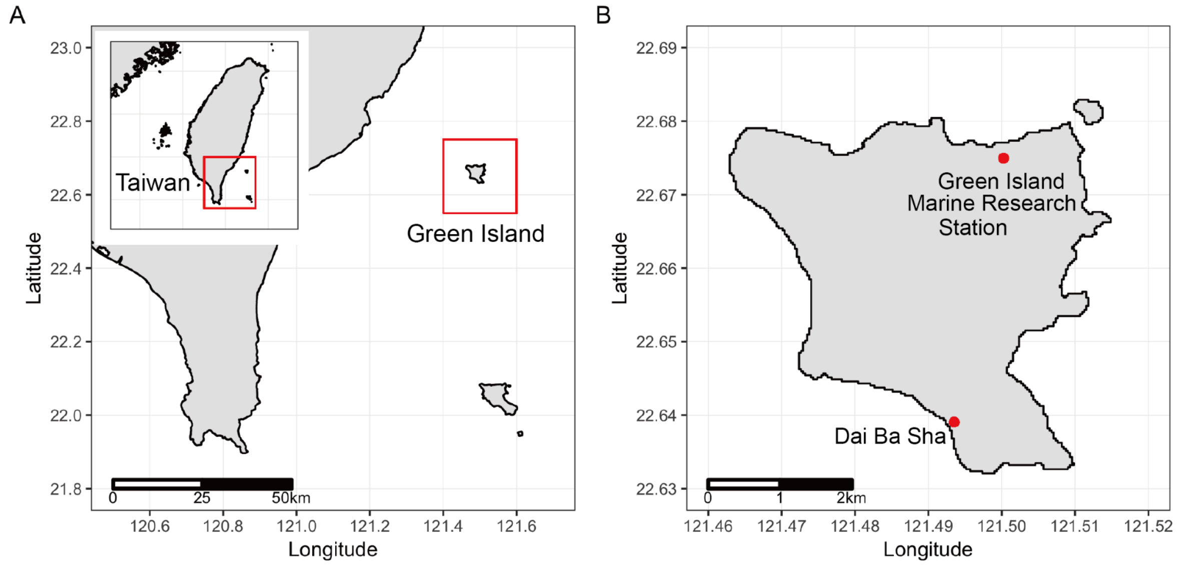


**Supplementary Figure 2:** Composition of single-base INDELs in homopolymers.

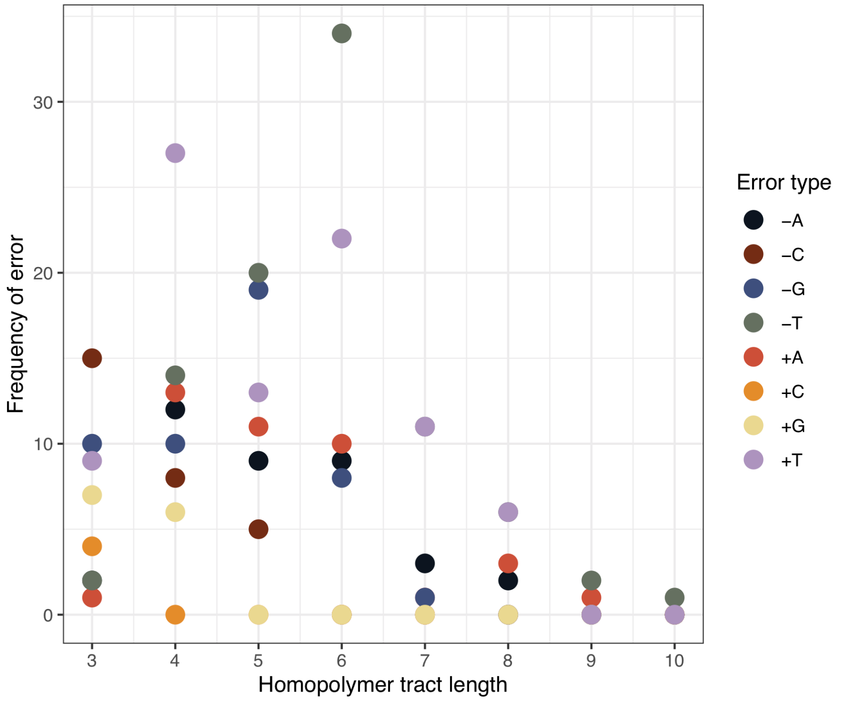

**Supplementary Figure 3**: ONT assembly feature of sample Ceb. (A) Dotplot against Illumina assembly. (B) AT content in 50 bp widows. (C) Nanopore and Illumina read coverage in 50 bp windows.

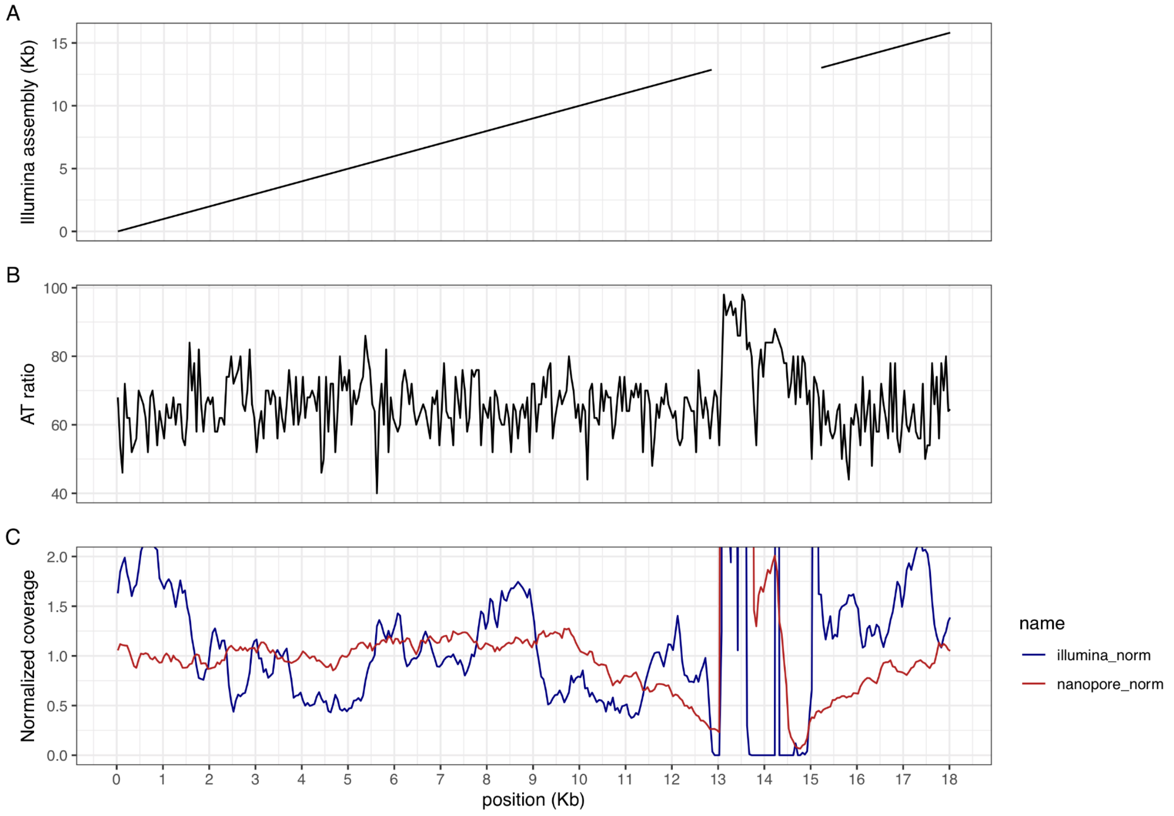

**Supplementary Figure 4**: Coalescence tree for Aplysiidae.

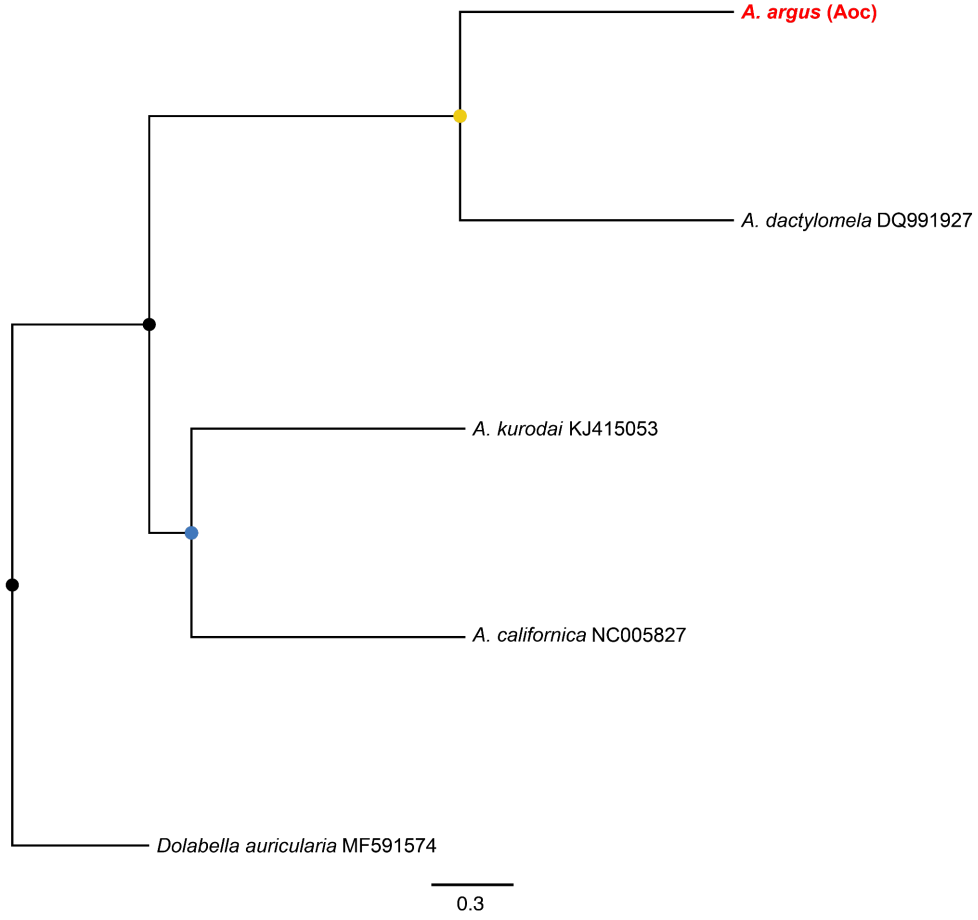

**Supplementary Figure 5**: Coalescence tree for Patellogastropoda.

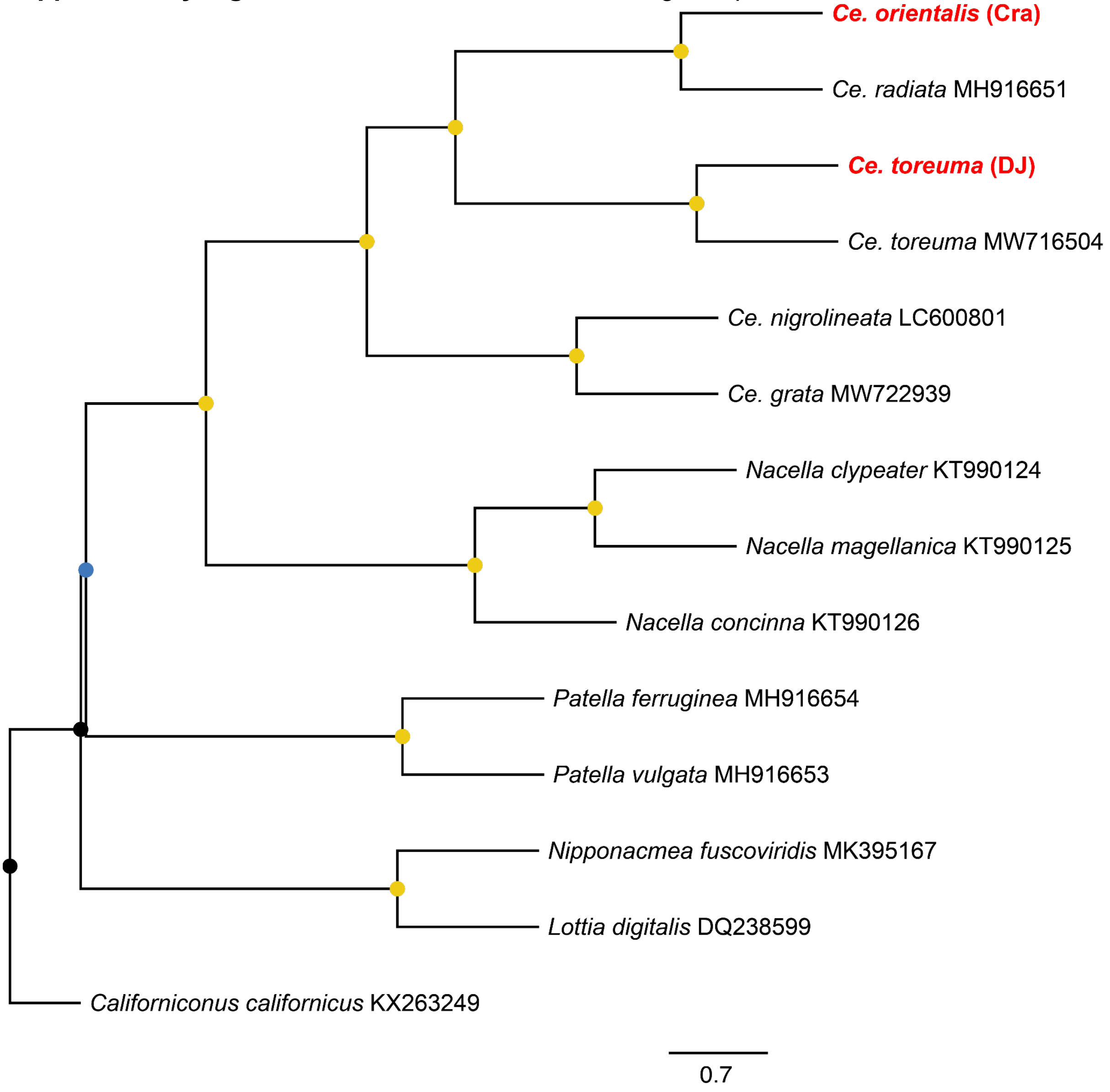

**Supplementary Figure 6**: Coalescence tree for Conidae.

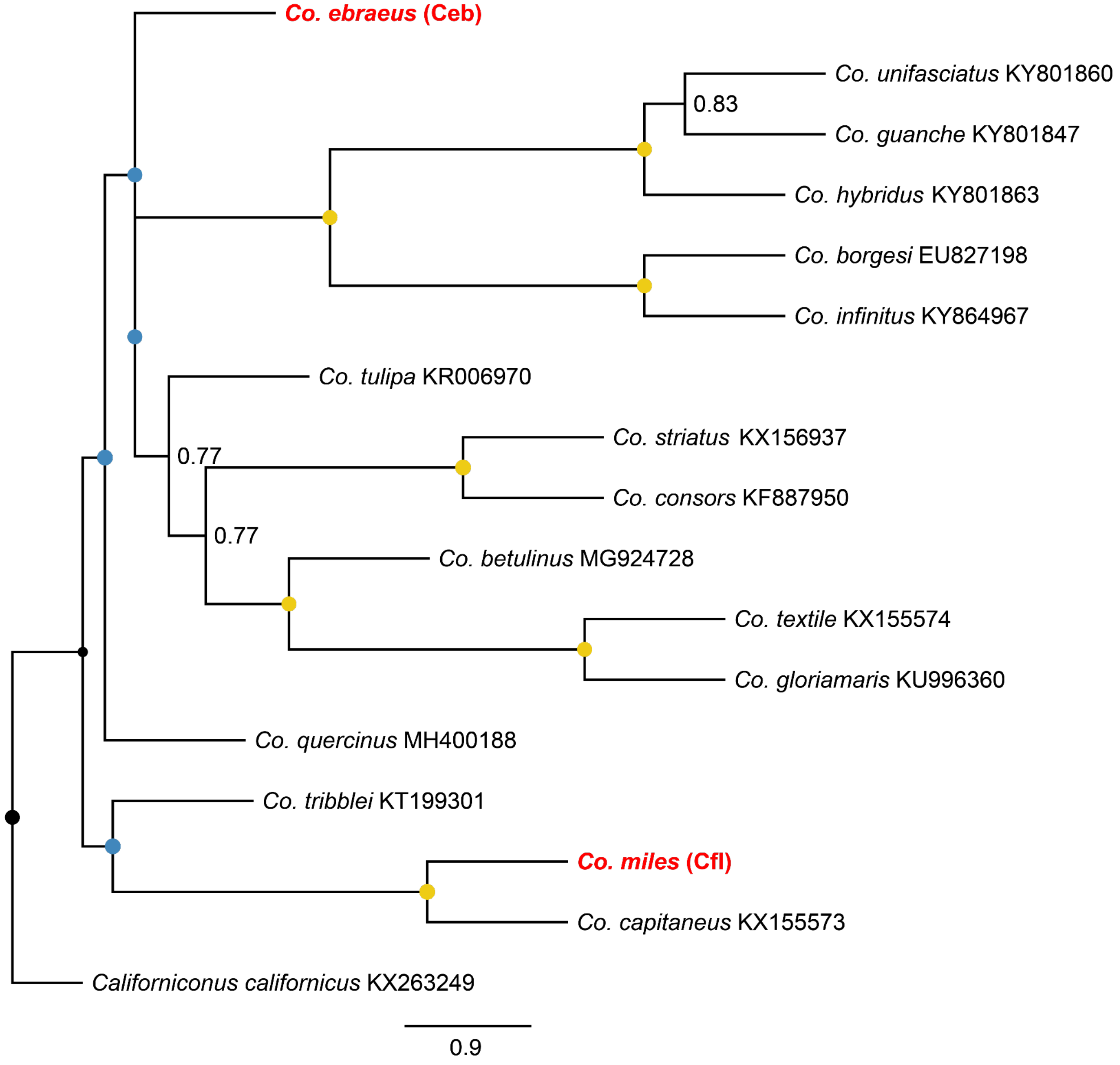

**Supplementary Figure 7**: Coalescence tree for Muricidae.

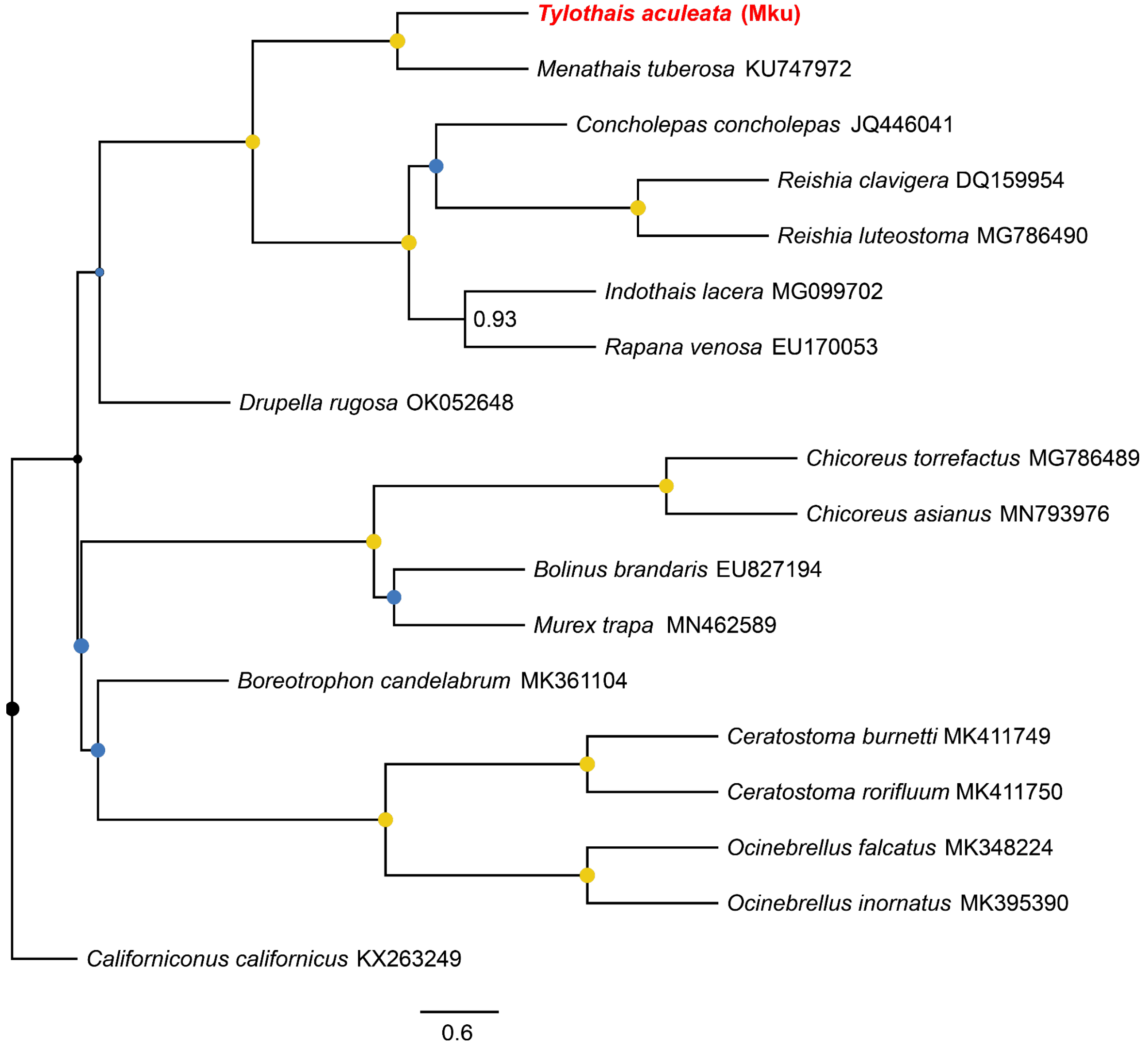

**Supplementary Figure 8**: Synteny plot for all the analysed mitogenomes used in this study.

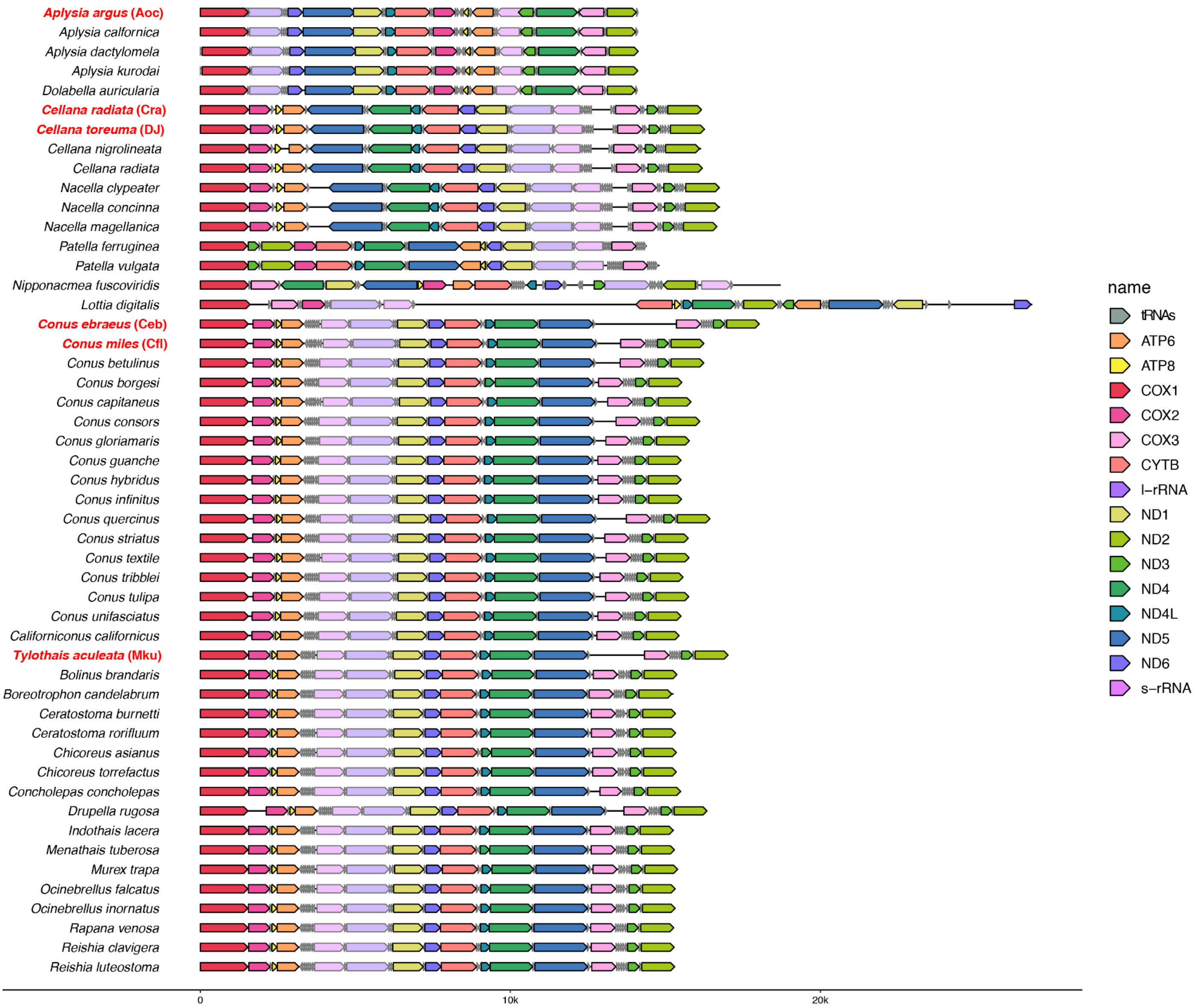
